# RPA and Rad27 limit templated and inverted insertions at DNA breaks

**DOI:** 10.1101/2024.03.07.583931

**Authors:** Yang Yu, Xin Wang, Jordan Fox, Qian Li, Yang Yu, P.J. Hastings, Kaifu Chen, Grzegorz Ira

**Author notes:** Equal contribution (Yang Yu, Xin Wang).

## Abstract

Formation of templated insertions at DNA double-strand breaks (DSBs) is very common in cancer cells. The mechanisms and enzymes regulating these events are largely unknown. Here, we investigated templated insertions in yeast at DSBs using amplicon sequencing across a repaired locus. We document very short (most ∼5-34 bp), templated inverted duplications at DSBs. They are generated through a foldback mechanism that utilizes microhomologies adjacent to the DSB. Enzymatic requirements suggest a hybrid mechanism wherein one end requires Polδ-mediated synthesis while the other end is captured by nonhomologous end joining (NHEJ). This process is exacerbated in mutants with low levels or mutated RPA (*rtt105*Δ; *rfa1*-t33) or extensive resection mutant (*sgs1*Δ *exo1*Δ). Templated insertions from various distant genomic locations also increase in these mutants as well as in *rad27*Δ and originate from fragile regions of the genome. Among complex insertions, common events are insertions of two sequences, originating from the same locus and with inverted orientation. We propose that these inversions are also formed by microhomology-mediated template switching. Taken together, we propose that a shortage of RPA typical in cancer cells is one possible factor stimulating the formation of templated insertions.

## INTRODUCTION

Templated insertions are a type of genome rearrangement wherein a segment of DNA is inserted into a chromosome, thereby duplicating existing genetic material. These types of copy number variations (CNVs) can lead to genetic disorders and are among the most common in cancer genomes (1,2). The inserted DNA may originate from nearby locations on the same chromosome (called hereafter local insertions) or from distant locations on the same or different chromosome (distant insertions). Templated insertions can be complex, involving multiple fragments copied from different parts of the genome (1,3). The size of templated insertions ranges from a few nucleotides to over a megabase. Multiple mechanisms have been proposed to explain templated insertions. Analysis of the junctions has demonstrated 0 to 20 nucleotides of microhomologies, implicating several DNA double-strand break repair pathways: nonhomologous end joining (NHEJ), alternative-end joining (Alt-EJ) and microhomology-mediated break-induced replication (MMBIR). In MMBIR, DSB ends can initiate low processivity, template switch-prone DNA synthesis at microhomologies (4,5). In a related model, one DSB end primes DNA synthesis while the other is captured by nonhomologous end joining (NHEJ), restoring chromosomal integrity (6). Support for such a hybrid mechanism comes from observations that DSB end mobility, Polδ, and NHEJ proteins are all necessary for at least some of the templated insertions at CRISPR/Cas9 induced events observed in cancer cells (6). However, a large fraction of the described events carry no microhomology on either DSB end, suggesting the involvement of a different polymerase than Polδ that requires initial microhomology, or that another mechanism contributes (6). Another proposed mechanism involves the capture of extrachromosomal DNA fragments at double-strand breaks (DSBs) by nonhomologous end joining (NHEJ) (3,7–9). The evidence supporting this mechanism includes microhomology typical for NHEJ at both ends of templated insertion and the requirement for key NHEJ enzymes. Various types of DNA can be captured at double-strand breaks (DSBs), including mitochondrial DNA, transposon cDNA, nuclear genomic DNA, as well as transformed single-stranded (ssDNA) or double-stranded DNA (dsDNA). Multiple fragments can be captured at DSBs, including a mixture of genomic DNA, mitochondrial DNA, and transposon DNA, as reported in yeast, humans, or plants (7,10–15). Alternative end joining (Alt-EJ) can result in very short local direct or inverted templated insertions, typically ranging from about 2 to 35 base pairs (16,17). These events require both PolQ and Polδ exonuclease activity to trim terminal nonhomology (18). Additionally, some models propose that specific inverted templated insertions can originate without any initial DSB but rather from template switches within replication forks followed by extrusion of inverted ssDNA, its replication and reintegration into the genome (19).

Although the analysis of cancer genome sequences has provided a wealth of information about templated insertions, the mechanisms leading to their formation can only be inferred (e.g. (1,20)). One experimental approach to model templated insertions and identify enzymes or conditions that upregulate these events is through the sequencing of double-strand break (DSB) repair products carrying rare templated insertions. Templated insertions have been observed at HO endonuclease or CRISPR/Cas9 induced DSBs in yeast and humans, as well as at programmed DSBs at the V(D)J locus (3,6,8,21). The vast majority of these events in yeast and humans are ∼50-300 base pairs in size. In wild-type yeast cells grown under optimal conditions, such insertions are very rare. However, in stressed stationary phase cells, they increase dramatically (8). The insertions from genomic DNA, mitochondrial DNA (mtDNA), or transposon cDNA increase by ∼100 to 2000 fold, with the mtDNA and cDNA being controlled by yeast EndoG, Nuc1 (8). Templated insertions also increase in cells deficient in the 5’ flap endonuclease and helicase, Dna2. The function of Dna2 in limiting such insertions is conserved from yeast to humans and can be related to its role in processing over-replicated DNA at reversed replication forks or long 5’ flaps (3,22–24). In support of this possibility, it was recently demonstrated that over-replicated DNA at reversed replication forks can recombine across the genome generating duplications (25). Genomic insertions observed in stationary phase cells or in *dna2*Δ mutants are not associated with deletion of the donor fragment, thus indicating a copy-paste mechanism. Inserted DNA in yeast and humans often originates from fragile regions of the genome, including telomeric and other repetitive regions, or R-loops (3,21). Microhomology-mediated templated insertions including local inversions have also been observed in other DSB inducible recombination assays. In these assays regular homology driven repair was interrupted and transiently switched to microhomology driven synthesis (e.g. (26–28)).

In this study, we investigated the role of several DNA repair/replication enzymes in the regulation of templated insertions at double-strand breaks (DSBs). Specifically, we focused on proteins previously shown to regulate microhomology-mediated annealing of single-stranded DNA (ssDNA), including RPA and the RPA regulatory factor Rtt105 (29–33). Deficiency in the RPA complex is commonly observed in cancer cells experiencing replication stress, and it leads to various types of genome instability (34–36). In this study, we also investigated the role of enzymes functionally related to Dna2: the 5’ flap endonuclease Rad27, which, similar to Dna2, processes 5’ flaps during replication, and the Sgs1 and Exo1 enzymes, which promote extensive resection along with Dna2 (24,37). RPA and *rad27*Δ mutants exhibited increased levels of templated insertions and displayed specific and distinct patterns of insertions. Furthermore, we characterized different mechanisms of local and distant *de novo* templated insertion with inverted orientation. Based on our findings in model system, we propose that a shortage of RPA in cancer cells is one possible factor stimulating the formation of microhomology-mediated templated insertions.

## MATERIAL AND METHODS

### Yeast strains

All yeast strains are JKM139 derivatives (38) (*MAT***a** DELho *hml*::*ADE1 hmr*::*ADE1 ade1 leu2-3,112 lys5 trp1*::*hisG ura3-52 ade3*::*GAL10*::*HO*) and are listed in **Supplementary Table S1**. All strains were constructed by transformation. The rfa1-t33 mutant was constructed using CRISPR/Cas9 by transforming the plasmid bYY37 carrying gRNA sequence (ATGGCAGCAACAGAACCTTC) with the plasmid bRA90 (a gift from James Haber) as the backbone and “rfa1-S373P” oligo (CCTATGGAATCAGCAAGCCCTTGATTTCAACCTTCCTGAAGGTCCAGTTGCTGCCATTAAAGGTGTTCGTGTGACGGATTTTG) as the template.

### Yeast growth and DSB induction

Yeast cells were grown overnight in YPD (1% yeast extract, 2% peptone, 2% dextrose), washed twice with YP-raffinose (1% yeast extract, 2% peptone, 2% raffinose) media, inoculated into 10 mL YP-raffinose media, and incubated at 30 °C overnight. When the cell density was about 2×10^7^/mL, 500 µL cells were spread onto YP-galactose (1% yeast extract, 2% peptone, 2% galactose, 2.5% agar) plates to induce HO expression and the DSB at *MAT***a** locus, and incubated at 30 °C for 5 days. Each sample was spread on 5-7 150×15 mm YP-galactose plates. The number of colonies was counted. All colonies on YP-galactose plates were collected, combined, and mixed vigorously. Approximately 80 μL cells were spun down for isolating genomic DNA.

### Amplicon sequencing of DSB repair products by *Break-Ins*

Amplicon sequencing was performed as previously described (8). Standard glass beads-phenol/chloroform/isoamyl alcohol (25:24:1) method was used for isolating genomic DNA. The DNA was dissolved in water and adjusted to a concentration of 10 ng/μL. The construction of the amplicon sequencing library was adapted from Illumina’s protocol #15044223 Rev. B. Briefly, the sequencing libraries were constructed with two rounds of PCR using KAPA HiFi HotStart ReadyMix polymerase (Roche, 7958927001). For the first round of PCR, 3.2 μL genomic DNA and 0.5 μL 10 μM primers were used for a total volume of 25 μL. The forward and reverse primers targeting the *MAT***a** locus are located 11 bp upstream or 18 bp downstream of the HO cleavage site respectively, containing Illumina adapter sequence and 3 bases unique home index as described (8). The thermocycling conditions were: 95 °C for 5 min; 22 cycles of 98 °C for 20 s, 65 °C for 30 s, 72 °C for 3 min; 72 °C for 10 min. After the first round of PCR, 18μL PCR product was purified with an equal volume of AMPure XP beads (Beckman Coulter, A63880). The DNA was eluted with 52.5 μL of 10 mM Tris (pH 8.5) and used as the templates for the second round of PCR. For the second round of PCR, 5 μL template DNA, 5 μL index primer N7xx and 5 μL index primer S5xx from Nextera XT V2 Index kit (Illumina, FC-131-2001) were used for the total volume of 50 μL. The following conditions were used for the second round of PCR: 95 °C for 5 min; 8 cycles of 95 °C for 30 s, 55 °C for 30 s, 72 °C for 3 min; 72 °C for 10 min. Then, the PCR product was purified with 1.2x volume of AMPure XP beads and eluted with 10 mM Tris pH 8.5. The average size of each sample was measured with TapeStation and the DNA concentration was measured with Qubit dsDNA BR Assay Kit (ThermoFisher, Q32850). Then, the DNA was diluted to a concentration of 4 nM. An equal volume of DNA from ∼20 samples was pooled into the library. The 0.2 M NaOH was used to denature the pooled library and PhiX library (Illumina, FC-110-3001). After diluting the libraries with pre-chilled HT1 to 12 pM, pooled library and PhiX library were mixed with the ratio 9:1, further denatured by incubating at 96 °C for 2 min and immediately put in an ice-water bath for 5 min. The MiSeq and Reagent Kit v3 (600 cycles) (Illumina, MS-102-3003) were used for sequencing. The cluster density for all libraries was ∼1100 K/mm^2^.

### Analysis of insertion of transformed DNA

The 69-mer reverse-phase cartridge-purified oligo ordered from Sigma-Aldrich were dissolved to 100 μM in TE buffer. The oligo CATTGAACAACATGTTGCTGTAAGNN NNACTCATA<u>GTACAGAC</u>GGCGTGTATCTGGTATTG<u>GTCTGTAC</u> contain 4-mer random nucleotides in the middle (NNNN), 8-nt inverted repeats (underlined) at the 3’ terminus with an 18-nt spacer. To analyze the insertions of transformed DNA, 10 mL cells were collected when the density reached ∼1.5 × 10^7^ cells/mL in YEP-Raffinose, washed twice with water and mixed with 240 μL 50% PEG3350, 36 μL 1M lithium acetate, 40 μL 100 μM DNA and 44 μL water to a total of 360 μL. After incubating at 30°C for 30 min followed by 42°C for 30 min, the cells were spun down, resuspended in 4.2 mL water, and spread on 150×15 mm YEP-Galactose plates (700 μL per plate). A total of 6 plates were spread. The plates were incubated at 30°C for 5 days. All colonies on YP-galactose plates were collected, pooled, and mixed vigorously. Approximately 80 μL cells were spun down for isolating genomic DNA for amplicon sequencing as above.

### Detection of insertions from Break-Ins analysis

The identification of insertions is outlined in the iDSBins protocol published previously (8) (https://github.com/gucascau/iDSBins.git). Briefly, we developed iDSBquality pipeline for raw reads preprocessing using BBDuk to eliminate Phix reads and to ensure reads quality control. Subsequently, the iDSBdetection pipeline was used, integrating PEAR and BLASTN alignments for the detection of reads featuring >10 bp insertions at DSB. Leveraging the information obtained from BLASTN alignments, we introduced iDSBdeduplication to effectively eliminate redundant reads. The identification of junction indels and microhomologies, as well as the donor number and annotation were accomplished through iDSBannotation. These tailored protocols collectively constitute a comprehensive approach for the systematic identification and characterization of large insertions at DSBs.

### Feature analysis of large insertion

The feature analysis of large insertions, including R-loop, ARS, telomere, tRNA, and tandem repeats, was performed using the package LargeInsertionFeature (https://github.com/gucascau/LargeInsertionFeature.git). Briefly, random control locations were generated by shuffling the real observed locations of the insertions using the algorithm BEDTools shuffle. Bedtools v2.253 was used to retrieve genomic features that overlapped with or were located close to the insertion donor sequences in the genome.

### Detection of local foldback *MAT*a insertions and distant inverted insertions

To identify local templated foldback inversions, we initially delineated the template-inverted sequence based on two inverted 10 bp long microhomologies next to the telomere-proximal DSB end. The seed sequence, consisting of the first 5 bp copied (TACTT, see Fig. 1B, first foldback inversion example) followed by a short *MAT***a** sequence corresponding to inverted microhomology (CAGTATA or CAGCATA), was used for scanning sequencing reads for foldback inversion. By constraining this seed sequence (TACTTCAGTATA or TACTTCAGCATA), we ensured selection of foldback inversions of 5 bp or longer. Subsequently, the scanning process extended across both PEAR merged reads and unmerged reversed reads. As occurrences were not detected in unmerged reads, our analysis exclusively focused on merged reads for subsequent analyses. Utilizing BWA mem, we mapped these reads against chromosome III, in which the *HML* and *HMR* regions have been masked. To accurately deduplicate *MAT***a** reads, analysis of CIGAR strings was performed, with special emphasis on avoiding sequencing errors at both sides of *MAT***a** regions. Insertions (I) or deletions (D) at either MAT-L (left junction) or MAT-R (right junction) regions due to sequencing errors were disregarded. Following this, we executed deduplication, measuring insertion size and read count supporting each *MAT***a** hotspot insertion. Duplicated reads were identified as those sharing identical sequences containing the inverted sequence and their upstream and downstream 5 bp. Irrespective of read number, each local foldback inversion sequence was counted as a single event. To determine the 2 bp foldback inversions, we specifically selected reads corresponding to this 2 bp, but not longer, inversions. To identify distant inversions, we screened the reads corresponding to complex insertions that contained two donor sequences from single locus but with reverse orientations.

**Figure 1.**
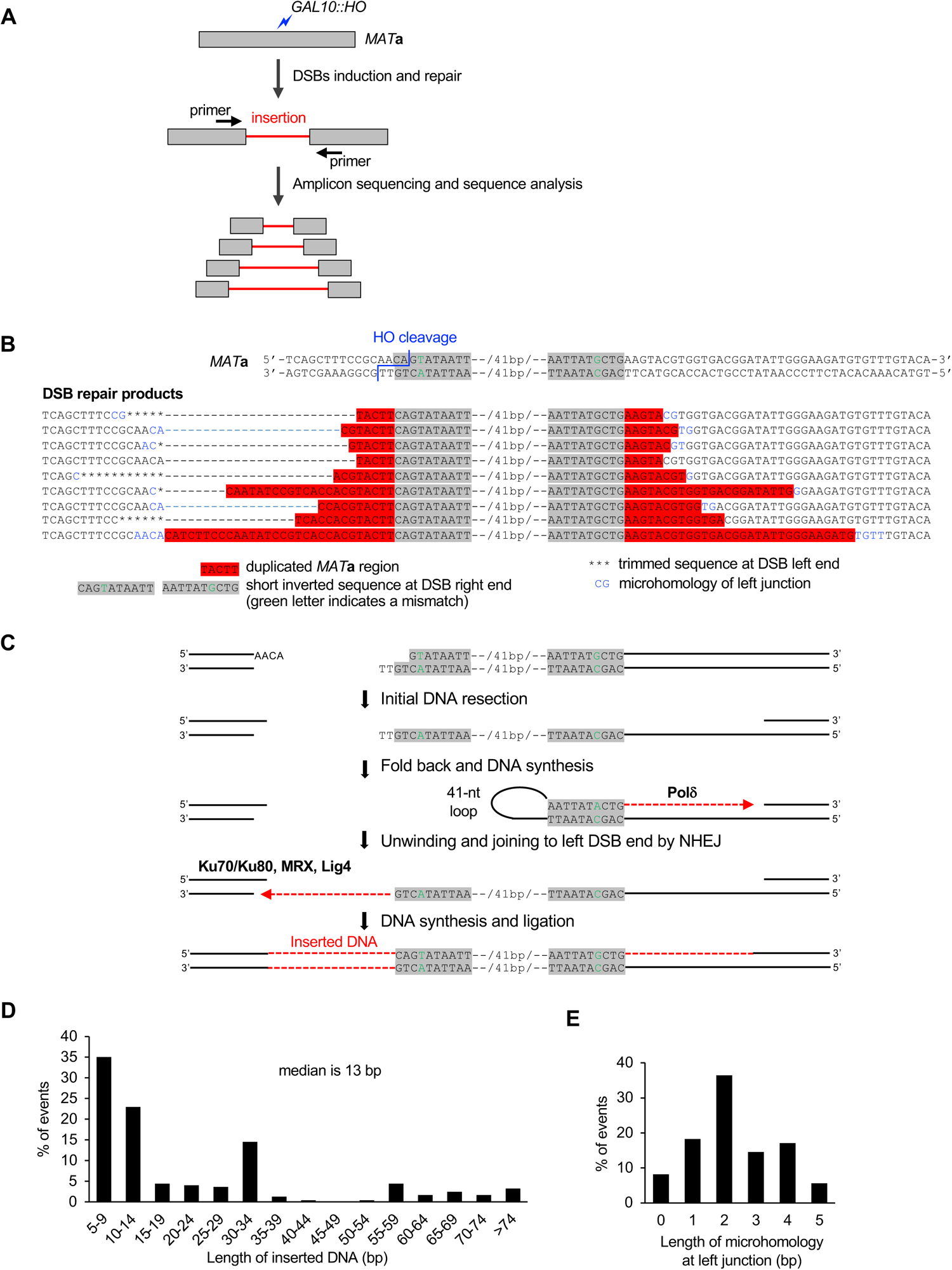
Local foldback inverted insertions at DSBs. (A) Scheme of the *Break-Ins* method for detecting insertions at DSBs. DSBs are induced by the HO endonuclease at the *MAT***a** locus. The DSB repair products are pooled and subjected to amplicon sequencing using the MiSeq platform. The analysis focuses on reads with >10 bp insertions at DSBs. (B) The inverted duplication from the region close to the HO cleavage site. Upper, the double strand *MAT***a** sequence including HO cleavage site. The HO induces DSBs (blue) with 4-nt 3’ overhangs. Lower, examples of the inverted insertions in DSB repair products copying the sequence (red) mediated by foldback 10-bp imperfect inverted microhomology sequences (grey) with a 41-nt spacer. The sequencing data of wild-type cells are taken from a previous publication (8). (C) Model presenting the mechanism of local foldback inversions at DSBs. The right end of HO-induced DSBs is resected to expose 10 nt imperfect inverted microhomologies (grey). The ssDNA then folds back to form a hairpin with 41 nt loop, and Polδ synthesizes a short DNA (red). The newly synthesized DNA unwinds from the template and ligates to the left DSB via NHEJ. The other strand is synthesized and ligated. (D) Size analysis of the foldback inversions from WT, *sgs1*Δ *exo1*Δ, *rfa1*-t33 and *rtt105*Δ mutants combined. Foldback inversions shorter than 5 bp were not included. (E) Length of microhomology at left junction from WT, *sgs1*Δ *exo1*Δ, *rfa1*-t33 and *rtt105*Δ mutants combined. Foldback inversions shorter than 5 bp were not included.

### Analysis of insertions originating from transformed ssDNA

Raw reads were pre-processed with quality control and merged with PEAR v0.9.11 as described above. Reads with any substring of transformed ssDNA (e.g., CATTGAACAA, CAACATGTTG, TTGCTGTAAG, ACTCATAGTA, GTGTATCTGG, TACCAGATAC, TACTATGAGT, or CTTACAGCAA) were recognized as insertions. Barcode sequences were defined between 5 bp upstream and 5 bp downstream of the barcode sequence. Perfect matches were required for these seed sequences. Thereafter, UMI features were used to distinguish between insertions with two identical and two different barcodes.

## RESULTS

### Formation of local foldback inversion at DSBs

To study templated insertions in yeast, we employed an HO endonuclease-inducible system where a single DSB per genome is induced at the *MAT***a** locus. This DSB cannot be repaired by homologous recombination because the strain lacks a homologous template, and both sister chromatids are cut by HO endonuclease. Instead, the DSB is repaired typically by NHEJ (38) or rarely by microhomology mediated repair such as Alt-EJ, altering the cleavage site. An amplicon sequencing-based method, *Break-Ins,* described previously, was used to sequence DSB repair products (8) (**Fig. 1A**). Analysis was limited to the sequencing reads that carry insertions of >10 bp at DSBs among all repair products. Previous analyses have focused on the transfer of mitochondrial DNA and transposon cDNA to the nucleus, as well as the function of Nuc1 in the degradation of these non-nuclear DNA species (8). Our aim here was to gain a better understanding of insertions originating from the nuclear genome. Analysis of over 100,000 independent DSB repair products in wild-type cells revealed local insertions originating from the right side of the double-strand break (DSB) that had identical microhomology at junctions, and varied in length from 10 to over 150 bp. All these insertions carried an inverted imperfect 10 bp microhomology. This inverted 10-bp sequence naturally occurs just 2 and 53 bp from the right DSB end and is separated by 41 bp (see **Fig. 1B**). We propose that resection of the right DSB end exposes two inverted repeats that anneal to each other forming a hairpin with a loop of 41 nt. After trimming of the terminal 2 nt at the 3’ end, new DNA synthesis is followed by displacement of the nascent strand that is then ligated to the second end of a DSB (**Fig. 1C**). We further screened the same sequencing reads for even shorter insertions from this locus and identified many more insertions of 5 to 9 bp (**Fig. 1B, 1D**). We note that identical sequencing reads corresponding to foldback inversion in single experiment were considered as single event; however, it is possible that multiple identical events occurred independently. Also, we cannot exclude PCR or sequencing bias and therefore some events are either underrepresented or not sequenced. Finally, it is possible that 1 to 4 bp insertions form by the same mechanism, but these are difficult to distinguish from other mechanisms during NHEJ. The left DSB end and the inverted sequence shared only 0-5 bp microhomology (median = 2 bp) indicating that the repair was completed by regular NHEJ (**Fig. 1E**). In conclusion, we observed very short foldback inversions formed at DSBs much shorter than the ones previously reported (e.g. (39)). Also, the mechanism of their formation is different, as explained below.

### NHEJ and Polδ are required for the formation of local foldback inversions

To investigate whether the formation of foldback inversions involves a hybrid mechanism, wherein one DSB end is extended by DNA synthesis at the hairpin, and the resulting duplicated region is unwound and captured by the second DSB end via nonhomologous end joining (NHEJ), we conducted an analysis of the genetic requirements for these events. First, we searched for the DNA polymerase involved in the DNA synthesis step. We found that elimination of Rev3 (Polζ) and Rad30 (Polη) had no effect on these events (**Fig. 2A, Supplementary Table S2**). However, elimination of Pol32, a component of Polζ and a nonessential component of Polδ needed for BIR (40), completely eliminated foldback inversions. Since *rev3*Δ showed no phenotype, we conclude that Polδ, rather than Polζ, is responsible for the extension of microhomology-mediated foldback structures. This finding is consistent with previous reports highlighting the important role of Polδ in stabilizing microhomologies and hairpin structures (28,31,41). Secondly, to assess the contribution of nonhomologous end joining (NHEJ) in foldback inversion events, we analyzed rare double-strand break (DSB) repair events in NHEJ-deficient mutants *yku70*Δ, *lig4*Δ, and *mre11*Δ. To increase the number of rare Alt-EJ events, ∼10-fold more cells were plated in these mutants. Our analysis revealed that Yku70, Lig4, or Mre11 were nearly essential for the occurrence of foldback inversions (**Fig. 2B**). Three remaining events likely resulted from alternative end joining (Alt-EJ) and exhibited 3 or 4 nucleotide microhomology at the double-strand break (DSB) end used to capture duplicated sequences. Overall, our findings support the view that both Polδ dependent DNA synthesis and nonhomologous end joining (NHEJ) are involved in the formation of very short foldback inversions.

**Figure 2.**
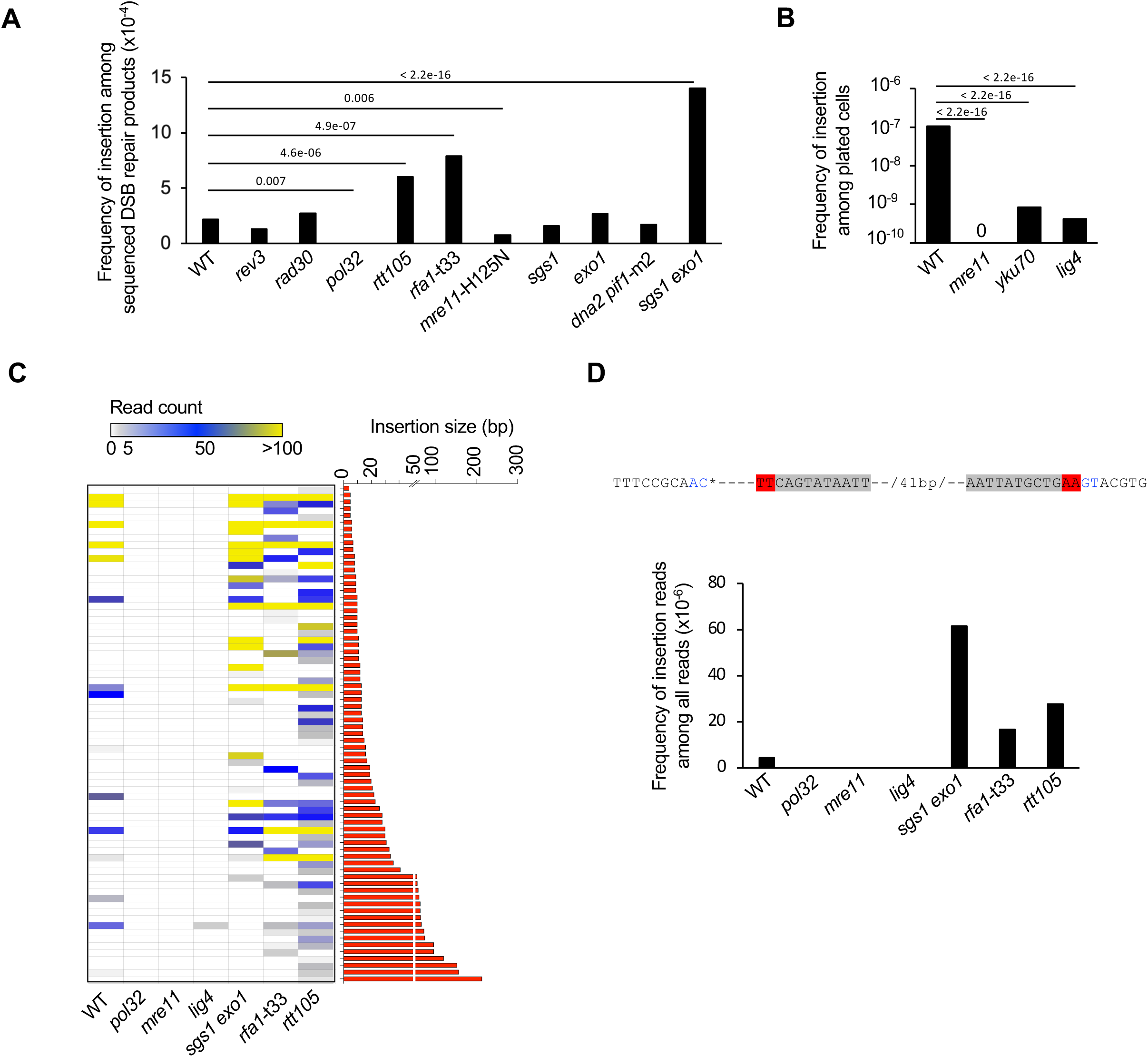
Genetic requirements for local foldback inversions at DSBs. (A) Analysis of foldback inversions in indicated mutants. Foldback inversions shorter than 5 bp were not included. P values were determined using χ^2^ test; n – number of DSB repair products tested for each mutant is shown in Table S2. (B) Foldback inversions in NHEJ mutants. P values were determined using χ^2^ test, n – number of cells plated is shown in Table S2. (C) Size and read number distribution for foldback inversions in indicated mutants. Each row represents a specific insertion event. Some of the events have the same size but different left junctions. (D) Number of reads corresponding to a 2-bp foldback inversion in indicated mutants. The upper is the sequence of inverted insertion (red), inverted microhomology is shown in grey.

### Rtt105, RPA and extensive resection suppress local foldback inversions

Next, we analyzed foldback events in RPA mutants, which were previously shown to affect microhomology-mediated repair (29–33). We tested the RPA mutant strain *rfa1*-t33, which exhibits decreased DNA binding, decreased growth, and defective replication (42–44) as well as the *rtt105*Δ mutant, which has a decreased level of nuclear RPA and reduced binding of RPA to single-stranded DNA (ssDNA) (33,45,46). The frequency of local foldback inversions mediated by microhomology among sequenced DSB repair products increased in both the *rtt105*Δ and *rfa1*-t33 mutant strains. (**Fig. 2A, 2C**). Previous studies have shown increased foldback inversions in the absence of Mre11 nuclease, which is known to process hairpin structures. Mre11 nuclease can process hairpin structures with short loops of up to approximately 8 nucleotides and inhibits foldback inversions (39,47). Hence, the 41-nucleotide loop observed here (**Fig. 1B-C**) is unlikely to be processed by Mre11. Indeed, we found no increase in foldback inversions in the *mre11*-H125N nuclease mutant, consistent with Mre11 having no role in the cleavage of this hairpin loop (**Fig. 2A**). Additionally, we investigated the role of extensive resection enzymes Sgs1, Dna2, and Exo1 in short foldback inversions. We observed an increase in local foldback inversions in the *sgs1*Δ *exo1*Δ mutant, while no change was seen in the single mutants *sgs1*Δ, *exo1*Δ, or *dna2*Δ *pif1-m2* (**Fig. 2A, 2C**). The MRX complex, along with Sae2, generates a few hundred nucleotides of ssDNA (37,48), which is sufficient to expose two inverted repeats located within the first 63 base pairs of the break. Without extensive resection, very short ssDNA intermediates of Mre11 mediated processing may limit RPA binding, which could further stimulate microhomology annealing.

In summary, the sequence and genetic analysis of foldback inversions described here supports a model in which the two ends of a single double-strand break (DSB) engage in two different pathways of repair. One end primes DNA synthesis at microhomology, while the other captures a duplicated region by nonhomologous end joining (NHEJ) (**Fig. 1C**). Our analysis focused on events with >5-nucleotide inversions, but it is highly likely that inversions as short as 1-4 base pairs are formed by a similar mechanism. These shorter inversions are challenging to distinguish from typical NHEJ errors associated with deletions. To further investigate, we compared the read number of the sequence corresponding to foldback-mediated synthesis of only 2 nucleotides (“TT”) in wild-type and mutant strains with higher or lower numbers of foldback inversions. We found that the read number was higher in *sgs1*Δ *exo1*Δ, *rfa1*-t33, or *rtt105*Δ mutants and absent in *pol32*Δ, *mre11*Δ or *lig4*Δ mutants. This suggests that the mechanism described here can lead to foldback inversions of just 2 nucleotides (**Fig. 2D**).

### Rad27, RPA and extensive resection suppress distant templated insertions

In addition to short local inverted templated insertions, we also observed distant templated insertions from all over the genome, as previously shown in Dna2 deficient cells (3,8). Besides the mutants tested above, we also looked in *rad27*Δ as it processes 5’ flaps during lagging strand synthesis like Dna2 (e.g. (24,49)). We also tested the Sgs1 and Exo1 enzymes, which, along with Dna2, promote extensive resection. Templated insertions from across the genome increased by approximately 10-30 folds in *rfa1*-t33, *rtt105*Δ, *rad27*Δ, and *sgs1*Δ *exo1*Δ mutants, but not in *sgs1*Δ, *exo1*Δ, or *mre11*-H125N mutants (**Fig. 3A-B, Supplementary Table S2**). Interestingly, Sgs1 and Rtt105 were found to suppress insertions from Ty cDNA, which might be related to their role in controlling Ty transposition (50,51). We have plotted insertions of Ty1 cDNA from *rtt105*Δ with highest number of cDNA insertions (n=98). As previously observed most insertions originate from 3’ LTR region, however pieces of nearly all Ty1 can be inserted (**Supplementary Fig. S1A**) (9,52).

**Figure 3.**
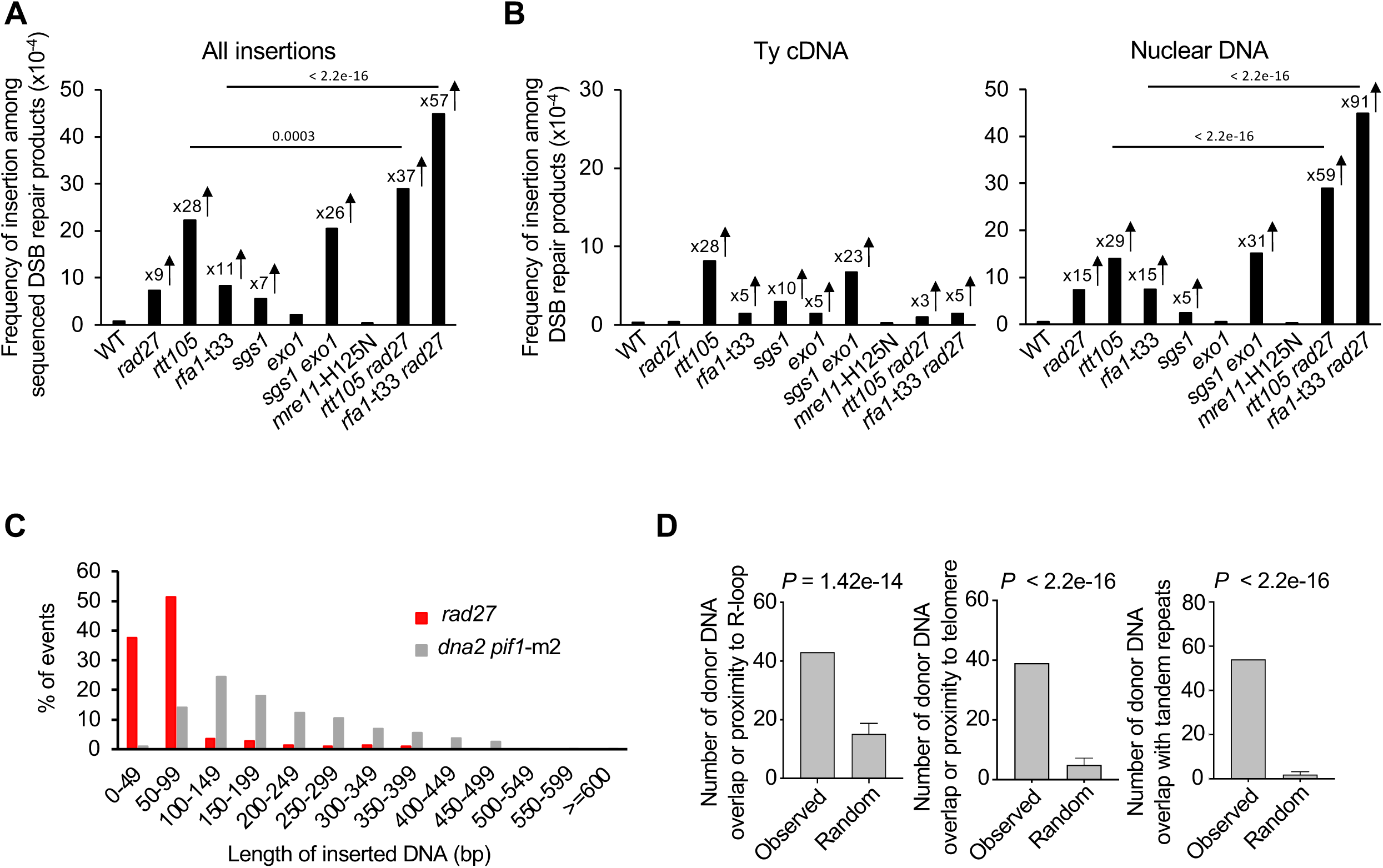
RPA and Rad27 suppress distant templated insertions. (**A, B**) Analysis of templated insertions in indicated mutants. Change in frequency of all insertions (**A**) and insertion of Ty cDNA or nuclear genome (**B**) among sequenced DSB repair products are shown. The insertions from *MAT***a** locus are excluded. P values were determined using χ^2^ test; n – number of DSB repair products tested for each mutant is shown in Table S2. (C) Size distribution of DNA inserted in *rad27Δ* mutant and *dna211 pif1*-m2 mutant. The data of *dna211 pif1*-m2 mutant are taken from a previous publication (8). Complex insertions were not included in the analysis. (D) Feature analysis of inserted DNA in *rad27Δ* mutant. P-values were calculated using a one-sided permutation test. Proximity is defined as a sequence within 0.2 kb from the R-loop, within 1 kb from telomere and the overlap with tandem repeats indicates at least 1 bp overlap.

The *rad27*Δ cells exhibited a 25-fold increase in insertions from the nuclear genome. Unlike in Dna2 deficient cells, very few insertions originated from rDNA (**Fig. 3B**, **Supplementary Table S2**). These insertions are templated (copy-paste and not deletion-paste) because we observed the expected number of insertions originating from essential genes (deletion within essential genes are expected to be lethal and are not detected by *Break-Ins*, see **Supplementary Fig. S1B**). Nearly all inserted DNA fragments are shorter than 100 bp, with a median size of 88 bp, and about 40% are shorter than 50 bp. This contrasts with *dna2*Δ mutants, in which most insertions (>85%) are longer than 100 bp, with a median size of 187 bp (**Fig. 3C**). The size of inserted DNA in *rad27*Δ corresponds to the size of 5’ flaps observed in *rad27*Δ cells by electron microscopy (24). Thus, alternative processing of 5’ flaps left in *rad27*Δ cells may be the source of fragments that are inserted at double-strand breaks. Analysis of 310 insertions in *rad27*Δ cells originating from throughout the genome shows that inserted DNA often overlaps with microsatellite DNA (1-12 nucleotides repeat) (**Fig. 3D**), or comes from within or next to telomeric repeats. Rad27 activity is particularly crucial for maintaining the stability of telomeric repeats and microsatellite DNA sequences (53–55) indicating that the inserted DNA might originate from loci most vulnerable to replication problems in *rad27*Δ. Three examples of the hotspots of the inserted DNA are shown (**see Supplementary Fig. S1C**). Inserted DNA also originated from previously mapped R-loops, consistent with the possible role of Rad27 in genome stability at R-loop generating loci (56).

We identified a combined total of 321 insertions originating from the nuclear genome in *rtt105*Δ and *rfa1*-t33 mutants, and these insertions also originated from fragile and repetitive sites of the genome (**see Supplementary Fig. S2**). We note that 45 insertions from *sgs1*Δ *exo1*Δ were not sufficient to establish the features of DNA inserted from the genome. Furthermore, the double mutants *rfa1*-t33 *rad27*Δ or *rtt105*Δ *rad27*Δ showed an additive effect on insertion level, suggesting that RPA and Rad27 operate in different pathways to prevent templated insertions at double-strand breaks. In conclusion, templated insertions are common in mutants defective in nucleases processing 5’ flaps, Rad27 or Dna2, as well as in RPA mutants.

### Distant inverted repeats are increased in *rad27*Δ and RPA mutants

Analysis of templated insertion events in *rad27*Δ, *rtt105*Δ, *rfa1*-t33, *dna2*Δ *pif1*-m2 mutants shows that 5-7% of them are complex, carrying at least two fragments (**Fig. 4A**). A frequent pattern of complex insertions was observed in *rad27*Δ (65.2%, 15/23), in which two fragments originate from the same locus but in inverted orientation (**Fig. 4A**). We note that complex insertions with two sequences from a single locus inserted in a direct orientation are rare. The inverted sequences had partially overlapped sequences with a median length of 50 bp and a distance of approximately 2-20 bp between microhomology junctions. We further found that similar double inverted insertions were also observed in *rtt105*Δ (78% of complex events, 14/18), *rfa1*-t33 (100%, 3/3), and less frequently in Dna2 deficient cells (6%, 9/144). However, in *rtt105*Δ mutants, the size of inverted sequences was longer (median = 150 bp), and the two inverted sequences from a single locus are rather adjacent to each other with little or no overlap (distance between junctions was between 30 and 119 bp). Interestingly, microhomologies between the entire complex insertion and DSB ends are typical for NHEJ (median = 2 bp, range of 0-4 bp), while microhomology between the two inverted sequences is longer (median = 6 bp, range of 2-11), suggesting a mechanism other than NHEJ generating the inversion itself (see **Fig. 4B-C**). An example of sequences and the structure of the double inverted insertion are shown (**Fig. 4D-E, Supplementary Fig. S3**). Taken together, we propose a model that involves both a microhomology-mediated DNA synthesis step to generate inversion and a NHEJ step that generates insertion (**Fig. 4F**). First, microhomology-mediated DNA synthesis occurs at short 5’ flaps (*rad27*Δ) or in the context of secondary structures formed in cells with limited RPA, leading to inversions. The longer microhomology at the junction between inverted fragments is consistent with possible involvement of microhomology-mediated DNA synthesis. Displacement of a newly synthesized strand with inverted sequences is followed by its insertion at the DSB and results in distant inverted insertion.

**Figure 4.**
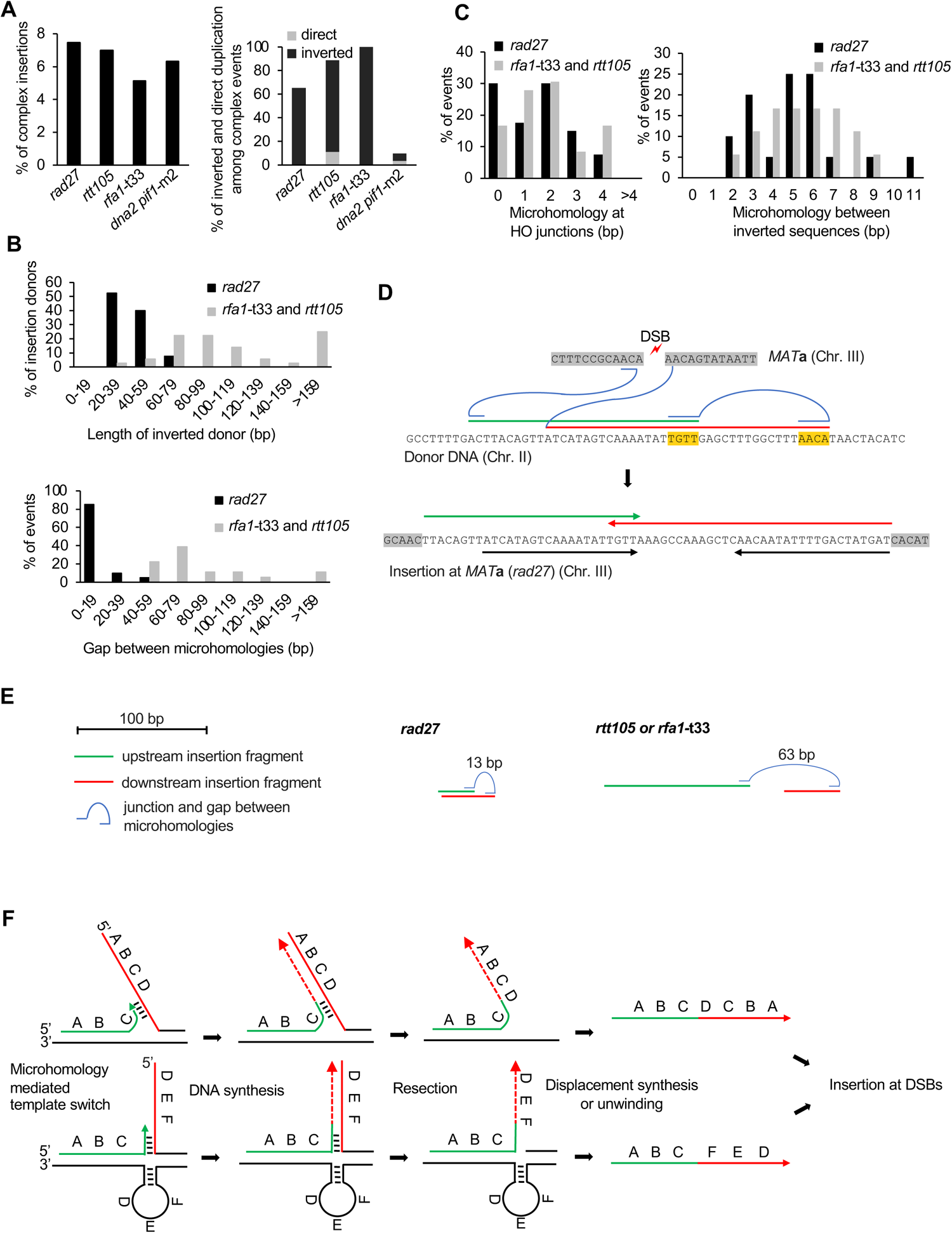
Distant inverted insertions in *rad27Δ* and RPA mutants. (A) Percentage of complex insertions among all insertions (left panel) and percentage of two sequence insertions from the same locus in inverted or direct orientation among complex insertions in indicated mutants (right panel, rDNA and Ty sequences are excluded). (B) Size analysis of insertion donor DNA (upper panel) and the size of the gap between two inverted microhomologous sequences (lower panel). (C) Microhomology analysis between inserted DNA and DSB ends (left) and microhomology between two inserted fragments (right). (D) An example of inverted insertion from *rad27*Δ cells. The upper panel shows the *MAT***a** sequence and two donor sequences (green and red) inserted at DSB. Blue curves mark the junctions, and the short horizontal lines represent the microhomology used for junctions. Highlighted in yellow is microhomology between inverted sequences. Final repair product carrying two inserted DNA fragments is shown below. The two black arrows represent inserted identical sequences with opposite orientations. (E) Scheme of the examples of inverted insertion events from *rad27*Δ and RPA mutants. The size of the gap between two inverted microhomologous sequences is indicated. (F) A model explaining distant inverted insertion at DSB. During replication in *rad27*Δ or RPA mutants, 5’ flaps or secondary structures can accumulate, respectively. Template switch at microhomologies initiates formation of inverted sequences. Upon resection and strand displacement, the inverted fragment is captured at DSB via NHEJ.

### Inversions can form within extrachromosomal pieces of ssDNA

To test directly whether such a mechanism is possible, we transformed single-stranded DNA (ssDNA) of 69 nt carrying an 8-nucleotide long internal inverted microhomology into wild-type cells followed by plating cells on YEP-Gal media to induce DSB at *MAT***a** locus (see **Fig. 5A**). Products of DSB repair with or without insertion of transformed fragments were collected and sequenced as described above. To distinguish between inversions generated by two molecules and those within a single molecule, we added random internal 4-nucleotide barcodes to the transformed DNA. Inversions formed within a single molecule would carry two identical barcodes, while inversions formed between two molecules would carry two different barcodes (see **Fig. 5A**), with four nucleotide barcodes providing 256 possible combinations.

**Figure 5.**
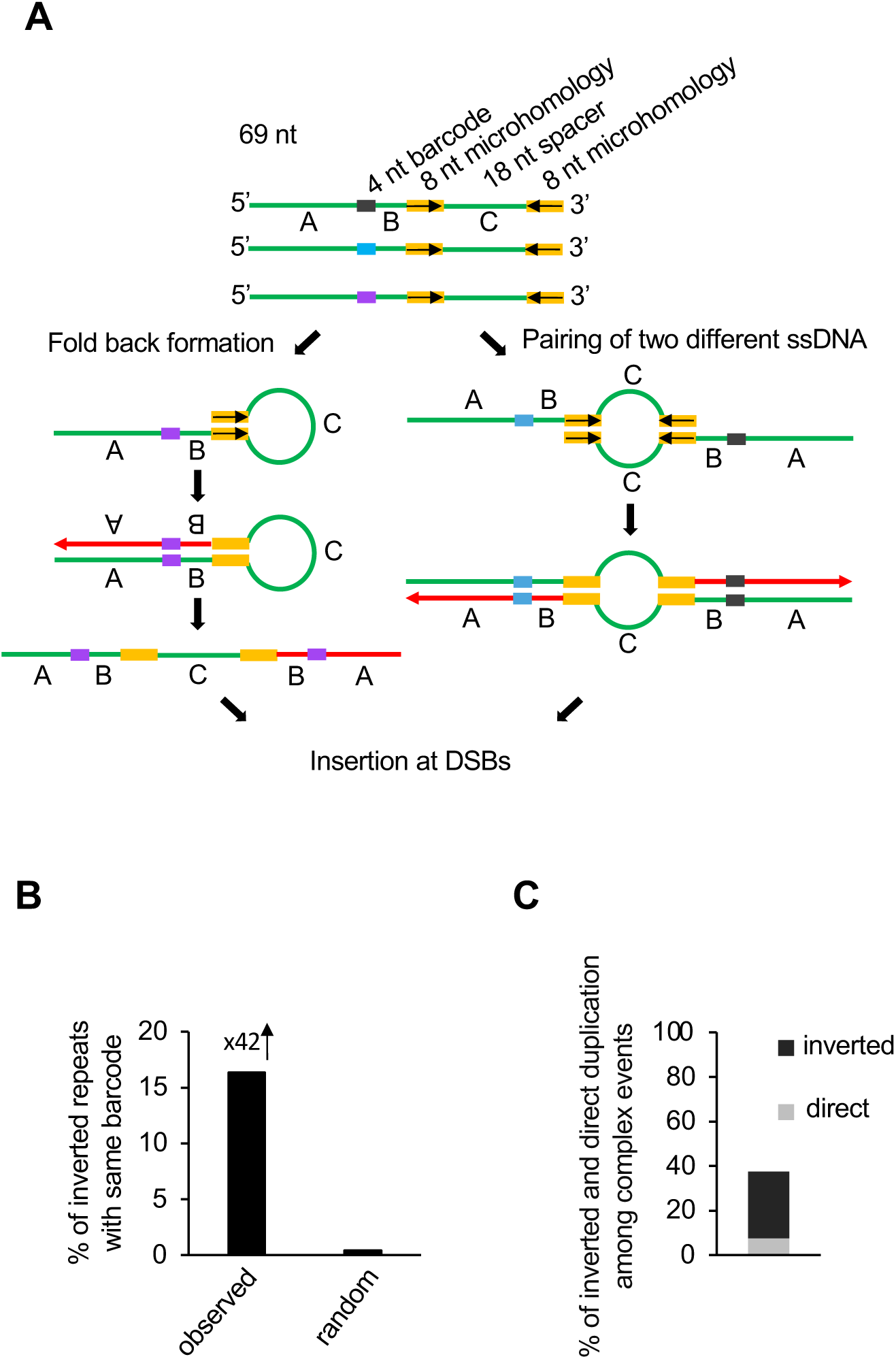
ssDNA can generate foldback inversions at DSBs. (A) The experimental system to test insertions of transformed ssDNA. The 69-nt ssDNA contains a 4-nt barcode in the middle, two 8-nt inverted microhomologies (yellow brackets), and an 18-nt spacer. The presence of barcodes within ssDNA allows differentiation between fold-back inversion (two identical barcodes) and pairing between two different ssDNA molecules (two different barcodes). (B) Bar graph shows random and observed percentage of inversions with identical barcode. (C) Number of complex insertions carrying two fragments from the same locus in inverted or direct orientation from wildtype stationary phase cells. The sequencing reads of wildtype stationary phase cells are taken from a previous publication (8).

Break-Ins analysis after the transformation of such ssDNA demonstrated a significant number of inverted insertions with barcodes (n=342), 16% of which were marked by identical barcodes. This is a much higher number than expected by chance (<1%) (see **Fig. 5B**). This result supports the possibility that inversions can occur within ssDNA, followed by its insertion at the double-strand breaks by NHEJ.

### Distant inverted repeats in stationary cells

We documented templated inverted insertions in mutants deficient in DNA replication and repair. To check whether similar events can occur in wild-type cells, we examined sequences of previously published templated insertions from stationary phase cells (8). In wild type stationary phase cells, templated insertions are far more common than in growing cells. While most of the insertions originate from mtDNA and Ty cDNA, stationary phase cells at 16 days exhibited ∼300-fold increase of templated insertions from all over the genome. About 12% of these insertions are complex events carrying two fragments, and among these events with two donors from nuclear genome (rDNA, Ty excluded), we found 30% carried sequences from the same locus and in inverted orientation. Therefore, distant inverted insertions can occur even in wild-type cells in suboptimal conditions (**Fig. 5C**).

## DISCUSSION

Templated insertions can occur at programmed breaks at the V(D)J locus or at endonuclease induced breaks (3,6,21,57). Despite their prevalence, the mechanisms underlying such events remain poorly understood, and the genetic pathways regulating these events are largely unknown. In this study, we utilized a yeast model organism and the *Break-Ins* method to sequence hundreds of thousands of double-strand break (DSB) repair products. Through this approach, we identified mutants with increased levels of simple and complex, local and distant templated insertions. Additionally, we have elucidated two mechanisms leading to the formation of either local or distant inversions.

In both wild type and most mutants tested, we observed local templated insertions that apparently proceed through a foldback inversion mechanism mediated by a short inverted microhomology located immediately next to the double-strand break (DSB). These foldback inversions ranged from 2 to approximately 150 base pairs (bp) in length, with the majority being under 14 bp. The requirements for Polδ, and NHEJ, as well as analysis of microhomologies at insertion junctions suggests a hybrid mechanism. One end primes DNA synthesis at local 10 bp microhomology while the other end uses NHEJ to complete repair. As a result, longer inverted repeats are formed (**Fig. 1C**). The first step of this mechanism shares some features typical for MMBIR: (i) synthesis is initiated at microhomology, (ii) newly synthesized DNA is unwound from the template, and (iii) a polymerase essential for break-induced replication (BIR) is responsible for the DNA synthesis step. In MMBIR, unwinding could lead to additional cycles of template switch and DNA synthesis (4), whereas here, the unwound strand is simply joined with the second end of the break by NHEJ or, rarely, by alternative end joining (Alt-EJ) (4). In humans, POLQ-mediated Alt-EJ is significantly more efficient than in yeast lacking POLQ, suggesting a potentially more prominent role in templated insertions. However, POLQ by itself predominantly generates very short templated insertions, typically up to ∼30 nucleotides in length (17,18). Moreover, studies have shown that POLQ is dispensable for templated insertions at CRISPR/Cas9 nuclease-induced DSBs (6). Comparable hybrid DSB repair to the one described here but initiated by homology driven strand invasion and DNA synthesis followed by second end capture by NHEJ was previously observed in experimental systems in human (e.g. (58)).

Local templated insertions were found to be limited by RPA, likely due to its known inhibitory role in the annealing of microhomologies. RPA plays a crucial role in preventing various types of microhomology-mediated genome instabilities (29–33). Lower levels of RPA also affected the occurrence of distant templated insertions from various fragile and repetitive regions of the genome. RPA plays a role in preventing secondary structure formation and regulates a broad range of DNA nucleases (e.g. (59–61)), activities that are likely important for preventing templated insertions. Cancer cells often experience a shortage of RPA due to replication stress and excessive firing of replication origins (34,35). Therefore, microhomology-mediated templated insertions in cancer cells could be partially attributed to insufficient RPA to protect all ssDNA. Extensive resection also facilitates foldback inversion, which could be related to lower RPA binding to very short ssDNA. Previous studies have shown that extensive resection is important to prevent *de novo* telomere formation at DSB, possibly by limiting binding of the Cdc13-Stn1-Ten1 complex (62,63), a complex structurally similar to RPA.

We have also observed an increased level of templated insertions from various genomic locations in *rad27*Δ cells. These insertions were shorter than those observed in any other mutant tested here and often contained microsatellite, telomeric DNA, and other repetitive regions. Rad27, the 5’ flap endonuclease (human FEN1), plays a crucial role in replication and stability at these loci (53–55,64,65). Thus, the pattern of templated insertions and the origin of inserted DNA closely align with the functions of Rad27. Notably, the most significant increase in templated insertions among mutants tested so far is observed in another 5’ flap endonuclease mutant, *dna2*Δ (3,8,22). DNA2 nuclease is commonly overexpressed in various cancer types, and this elevated level of DNA2 is often essential for the survival of cancer cells (66–69). This suggests that the number of substrates processed by DNA2 increases in cancer cells, and the high number of templated insertions observed could be related to these substrates.

In RPA mutants (*rtt105*Δ or *rfa1*-t33) and in *rad27*Δ mutants, we observed a common pattern of complex templated insertions consisting of two sequences from the same locus but in inverted orientation. Notably, in *rad27*Δ mutants the inverted sequences overlap with each other thus forming inverted repeats, whereas in RPA mutants they mostly do not overlap. Analysis of previously published complex templated insertions from Dna2 deficient cells or even from wild-type stationary phase cells also showed cases of both overlapping and non-overlapping inverted sequences. We propose that such inversions arise through microhomology-mediated template switch within single-stranded DNA intermediates of replication, followed by displacement of inverted sequence and its insertion at the DSB (model depicted in **Fig. 4**). Alternatively, inversions can occur within extrachromosomal DNA. Supporting this possibility, extrachromosomal transformed single-stranded DNA (ssDNA) carrying internal inverted microhomologies forms such inversions at DSBs with high frequency. The presence of longer microhomology between two inverted sequences supports the possibility that DNA synthesis was involved in its formation. Recently, the formation of extrachromosomal DNA with inverted DNA that reintegrates into the genome was postulated to explain the formation of inversions (19). Similar to the model proposed here, inverted sequences form through template switching at microhomologies during replication followed by strand displacement. It will be of great interest to investigate whether distant inverted templated insertions, as described here, can also occur in human cancer genomes. The high level of inverted microhomologies suggests that such events can occur nearly anywhere in human or other genomes (70,71). Once formed, overlapping inverted sequences (inverted repeats) can give rise to secondary structures within the genome, thereby promoting additional genome instabilities (39,70,72–82). Altogether, this work provides insights into origin of templated insertions that can be tested in cancer cells in future.

## DATA AVAILABILITY

The *Break-Ins* data has been archived at GEO under the accession number GSE260753. All original code has been deposited on GitHub. For local inversion insertion from *MAT***a**, the corresponding code is available at: https://github.com/gucascau/iMMBIR.git. The codes utilized for characterizing inverted repeats resulting from ssDNA transformation have been made available at: https://github.com/gucascau/iDSBInvert.git. The code for large insertion detection can be accessed at the following GitHub repository: https://github.com/gucascau/iDSBins.git. The features related to large insertion analysis have been uploaded to a dedicated GitHub repository: https://github.com/gucascau/LargeInsertionFeature.git. Any additional information required to reanalyze the data reported in this paper is available from the lead contact upon request.

## SUPPLEMENTARY DATA

Supplementary Data are available at NAR online.

## AUTHOR CONTRIBUTIONS

Y.Y. (BCM) constructed strains and conducted most of the experiments. X. W. and K. C. analyzed sequencing data and conducted most of statistical data integration; Y.Y. (BCH) assisted with software development and data analysis. J.F. and Q.L. performed some of the *Break-Ins* analyses; P.H. helped with data interpretation; G.I., Y.Y. (BCM), X. W., and K.C. designed experiments, analyzed the data, and wrote the manuscript. J.F. edited the manuscript.

## ACKNOWLEDGEMENTS

We thank Dr. James Haber and Dr. Anna Malkova for critical reading of the manuscript.

## Supplementary Figure Legends

**Supplementary Figure 1.**
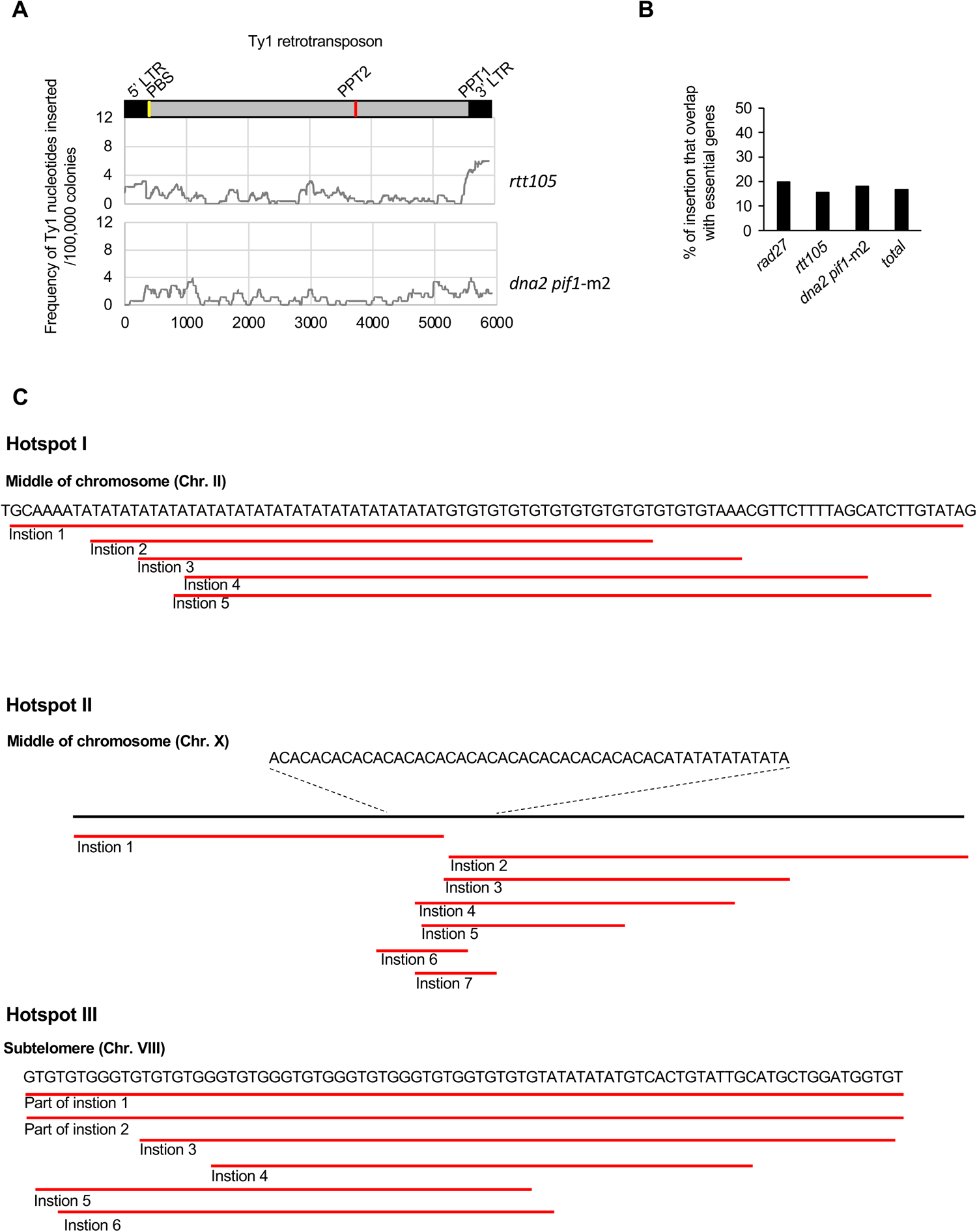
Analysis of inserted DNA. (A) Analysis of Ty1 sequences inserted at DSBs. Frequency of each Ty1 nucleotide insertion per 10^6^ of DSB repair products in *rtt105*Δ and *dna2*Δ *pif1*-m2 mutant cells. Schematic of Ty1 shown at the top indicating long terminal repeats (LTR), polypurine tracts (PPT1/2), and primer binding site (PBS). (B) The percentage of insertion overlapping with essential genes in indicated mutants and the percentage of essential genes in yeast genome (total). (C) The hotspots of inserted DNA from repetitive regions of the genome. The red lines indicate different inserted sequences from the same locus.

**Supplementary Figure 2.**
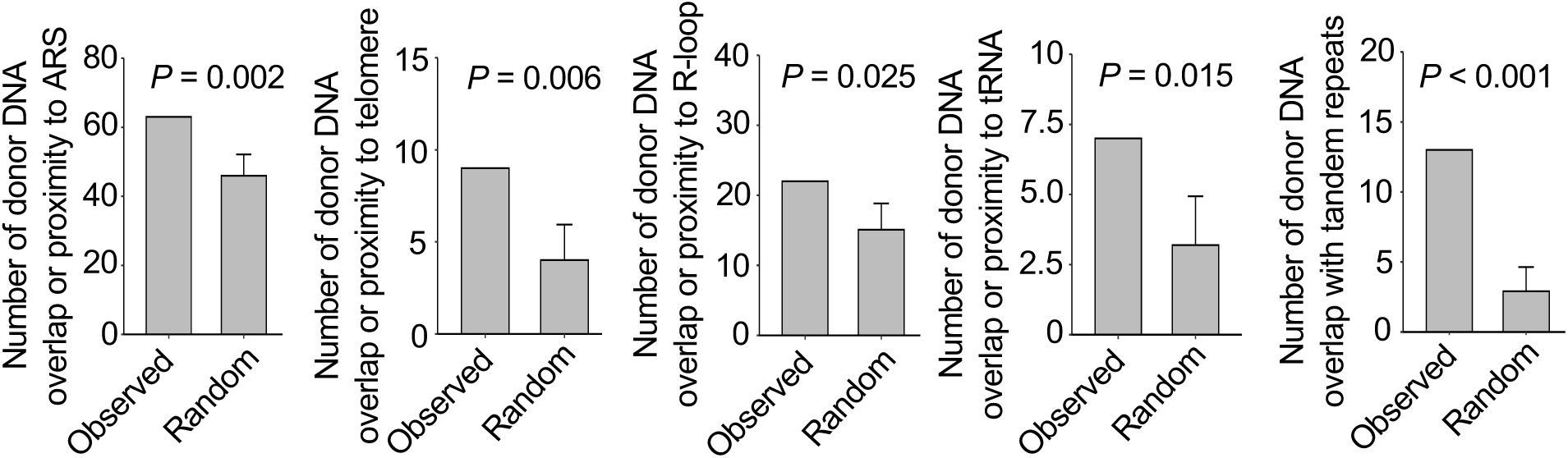
Feature analysis of inserted DNA in *rtt105*Δ mutant and *rfa1*-t33 mutants. The data of *rtt105*Δ mutant and *rfa1*-t33 mutant was combined for this analysis. P-values were calculated using a one-sided permutation test. Proximity is defined as a sequence within 0.2 kb from the R-loop or tRNA, within 1 kb from ARS or telomere, and the overlap with tandem repeats indicates at least 1 bp overlap.

**Supplementary Figure 3.**
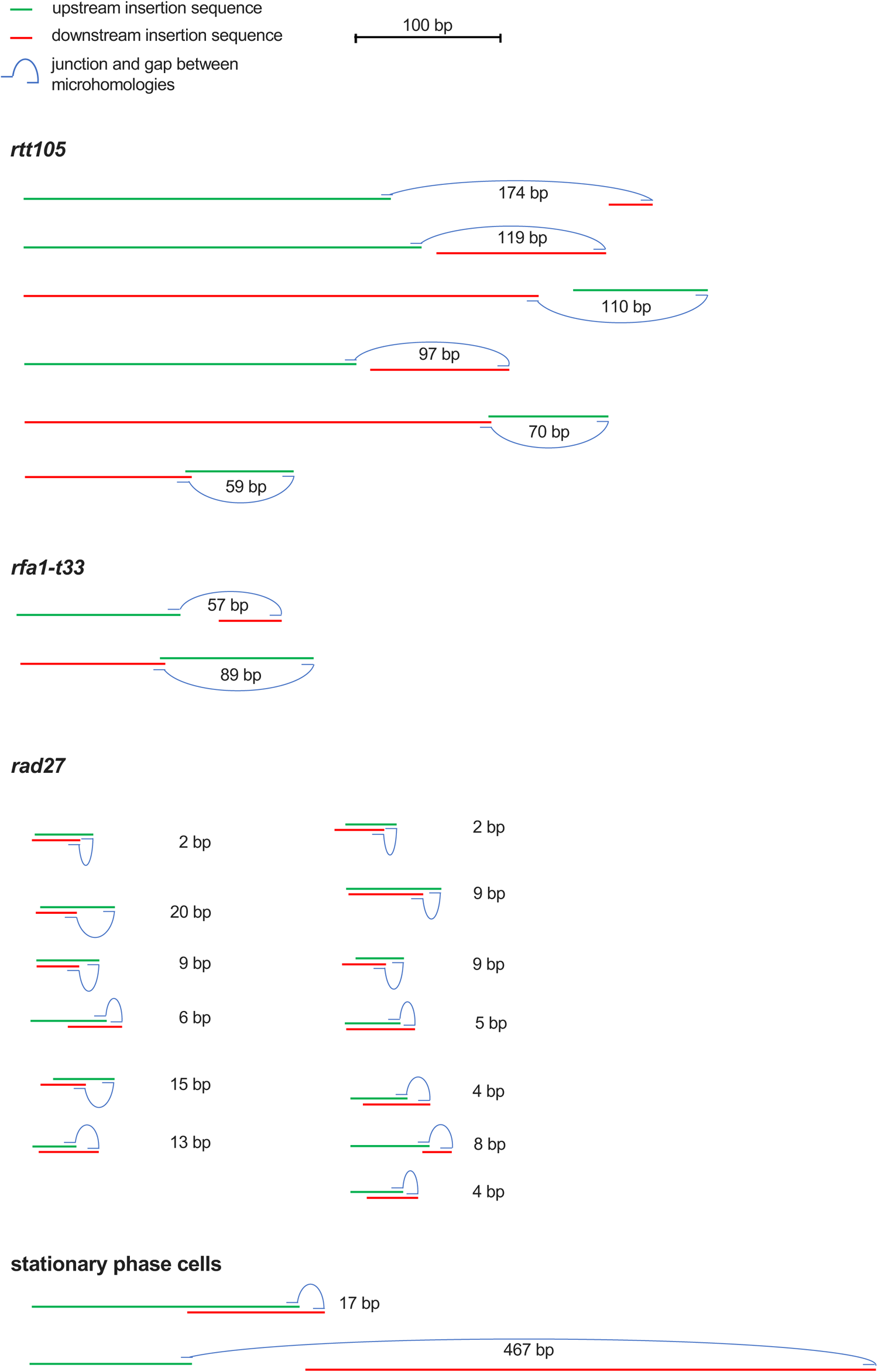
Scheme of the examples of inverted insertion events from *rad27*Δ, *rtt105*Δ, *rfa1*-t33 mutant and wildtype stationary phase cells. The size of the gap between two inverted microhomologous sequences is marked.

**Table S1.** List of strains used in this study.

| Strain name | Parental strain | Genotype | Source |
| --- | --- | --- | --- |
| JKM139 |  | DELho *hml::ADE1 MATa hmr::ADE1 ade1 leu2-3,112 lys5 trp1::hisG ura3-52 ade3::GAL10::HO* | (1) |
| yYY398 | JKM139 | *rad27::klTRP1* | (2) |
| yYY591 | JKM139 | *rtt105::hphMX* | this study |
| yYY616 | JKM139 | *rfa1-t33* | this study |
| yYY596 | JKM139 | *rtt105::hphMX rad27::klTRP1* | this study |
| yYY643 | JKM139 | *rfa1-t33 rad27::klTRP1* | this study |
| yWH475 | JKM139 | *dna2::kanMX pif1-m2* | (3) |
| yGI199 | JKM139 | *sgs1::KanMX exo1::TRP1* | (3) |
| yGI200 | JKM139 | *sgs1::KanMX* | (3) |
| yGI198 | JKM139 | *exo1::TRP1* | (3) |
| yZZ540 | JKM139 | *mre11-H125N::URA3* | (4) |
| yYY592 | JKM139 | *mre11::hphMX* | this study |
| yYY361 | JKM139 | *rev3::klTRP1* | this study |
| yYY590 | JKM139 | *rad30::kanMX* | this study |
| yYY387 | JKM139 | *pol32::natMX* | (5) |
| yYY119 | JKM139 | *yku70::TRP1* | this study |
| yYY399 | JKM139 | *lig4::klTRP1* | (5) |

1. Moore, J.K. and Haber, J.E. (1996) Cell cycle and genetic requirements of two pathways of nonhomologous end-joining repair of double-strand breaks in *Saccharomyces cerevisiae*. *Mol Cell Biol*, **16**, 2164-2173. 2. Yu, Y., Pham, N., Xia, B., Papusha, A., Wang, G., Yan, Z., Peng, G., Chen, K. and Ira, G. (2018) Dna2 nuclease deficiency results in large and complex DNA insertions at chromosomal breaks. *Nature*, **564**, 287-290. 3. Zhu, Z., Chung, W.H., Shim, E.Y., Lee, S.E. and Ira, G. (2008) Sgs1 helicase and two nucleases dna2 and exo1 resect DNA double-strand break ends. *Cell*, **134**, 981-994. 4. Shim, E.Y., Chung, W.H., Nicolette, M.L., Zhang, Y., Davis, M., Zhu, Z., Paull, T.T., Ira, G. and Lee, S.E. (2010) *Saccharomyces cerevisiae* Mre11/Rad50/Xrs2 and Ku proteins regulate association of Exo1 and Dna2 with DNA breaks. *EMBO J*, **29**, 3370-3380. 5. Yu, Y., Wang, X., Fox, J., Yu, R., Thakre, P., McCauley, B., Nikoloutsos, N., Li, Q., Hastings, P.J., Dang, W. *et al.* (2023) Yeast EndoG prevents genome instability by degrading cytoplasmic DNA. *bioRxiv*.

**Supplementary Table 2.** Frequency and types of DNA insertions.

|  |  |  |  |  |  |  | donor |  |  |  |  |  | frequency per 10<sup>4</sup> DSB repair products |  |  |  |  |  |  |
| --- | --- | --- | --- | --- | --- | --- | --- | --- | --- | --- | --- | --- | --- | --- | --- | --- | --- | --- | --- |
| strain name | genotype | number of colonies | number of colonies normalized by HO cut (n) | number of cells plated | all insertions excluding *MATa* | complex insertions | *MATa* * | mtDNA | Ty DNA | all nuclear DNA | rDNA | nuclear w/o rDNA | all insertions excluding *MATa* | *MATa* * | mtDNA | Ty DNA | all nuclear DNA | rDNA | nuclear w/o rDNA |
| JKM139 | WT** | 102976 | 101711 | 210000000 | 8 | 0 | 22 | 0 | 3 | 5 | 2 | 3 | 0.8 | 2.2 | 0.0 | 0.3 | 0.5 | 0.2 | 0.3 |
| yYY398 | *rad27* | 600672 | 445695 | 730000000 | 323 | 25 | 17 | 2 | 18 | 326 | 16 | 310 | 7.2 | 0.4 | 0.0 | 0.4 | 7.3 | 0.4 | 7.0 |
| yYY591 | *rtt105* | 374304 | 251798 | 720000000 | 560 | 40 | 151 | 42 | 205 | 353 | 63 | 290 | 22.2 | 6.0 | 1.7 | 8.1 | 14.0 | 2.5 | 11.5 |
| yYY616 | *rfa1*-t33 | 53184 | 48257 | 360000000 | 40 | 3 | 38 | 0 | 7 | 36 | 5 | 31 | 8.3 | 7.9 | 0.0 | 1.5 | 7.5 | 1.0 | 6.4 |
| yYY596 | *rtt105 rad27* | 111840 | 103358 | 360000000 | 299 | 11 | 27 | 1 | 10 | 299 | 16 | 283 | 28.9 | 2.6 | 0.1 | 1.0 | 28.9 | 1.5 | 27.4 |
| yYY643 | *rfa1*-t33 *rad27* | 14976 | 14254 | 240000000 | 64 | 3 | 6 | 1 | 2 | 64 | 2 | 62 | 44.9 | 4.2 | 0.7 | 1.4 | 44.9 | 1.4 | 43.5 |
| yWH475 | *dna2 pif1*-m2** | 197264 | 178507 | 1120000000 | 3966 | 250 | 31 | 0 | 255 | 3963 | 625 | 3338 | 222.2 | 1.7 | 0.0 | 14.3 | 200.9 | 31.7 | 187.0 |
| yGI199 | *sgs1 exo1* | 55000 | 37156 | 440000000 | 76 | 4 | 52 | 0 | 25 | 56 | 11 | 45 | 20.5 | 14.0 | 0.0 | 6.7 | 15.1 | 3.0 | 12.1 |
| yGI200 | *sgs1* | 106080 | 75160 | 200000000 | 42 | 1 | 12 | 3 | 22 | 18 | 3 | 15 | 5.6 | 1.6 | 0.4 | 2.9 | 2.4 | 0.4 | 2.0 |
| yGI198 | *exo1* | 54240 | 41429 | 120000000 | 9 | 0 | 11 | 1 | 6 | 2 | 0 | 2 | 2.2 | 2.7 | 0.2 | 1.4 | 0.5 | 0.0 | 0.5 |
| yZZ540 | *mre11*-H125N | 144320 | 142941 | 200000000 | 6 | 1 | 11 | 0 | 3 | 4 | 1 | 3 | 0.4 | 0.8 | 0.0 | 0.2 | 0.3 | 0.1 | 0.2 |
| yYY361 | *rev3* | 64160 | 60274 | 200000000 | 4 | 0 | 8 | 0 | 3 | 1 | 0 | 1 | 0.7 | 1.3 | 0.0 | 0.5 | 0.2 | 0.0 | 0.2 |
| yYY590 | *rad30* | 37296 | 32924 | 120000000 | 0 | 0 | 9 | 0 | 0 | 0 | 0 | 0 | 0.0 | 2.7 | 0.0 | 0.0 | 0.0 | 0.0 | 0.0 |
| yYY387 | *pol32* | 44160 | 39989 | 120000000 | 2 | 0 | 0 | 0 | 1 | 1 | 0 | 1 | 0.5 | 0.0 | 0.0 | 0.3 | 0.3 | 0.0 | 0.3 |
| yYY592 | *mre11* | 52176 | 10892 | 1800000000 | 0 | 0 | 0 | 0 | 0 | 0 | 0 | 0 | 0 | 0*** | 0 | 0 | 0 | 0 | 0 |
| yYY119 | *yku70* | 46960 | 1025 | 2400000000 | 0 | 0 | 2 | 0 | 0 | 0 | 0 | 0 | 0 | 8.3E-10*** | 0 | 0 | 0 | 0 | 0 |
| yYY399 | *lig4* | 65280 | 1223 | 2400000000 | 0 | 0 | 1 | 0 | 0 | 0 | 0 | 0 | 0 | 4.2E-10*** | 0 | 0 | 0 | 0 | 0 |

\* indicates the local foldback inversions from *MATa* locus related to Figure 1 and Figure 2
\*\* indicates data from Yu Y *et al* (8)
\*\*\* indicates the frequency per cells plated

